# Spontaneous drumming behaviour in a Galah

**DOI:** 10.64898/2026.03.25.714111

**Authors:** Joshua S. Bamford, Amanda R. Bamford

## Abstract

Drumming—rhythmic, percussive sound production using body parts or external objects—is rare among non-human animals, with confirmed tool-assisted cases previously limited to primates and Palm Cockatoos. Here, we report the first documented instance of spontaneous, tool-assisted drumming in a Galah (*Eolophus roseicapilla*). A captive, male Galah produced rhythmic tapping by striking a coconut shell against a metal bowl. Across 14 recorded sessions, the bird displayed consistent temporal structure characterised by two stable tapping rates (approximately 0.8 s and 0.2 s inter-onset intervals) arranged into recurring phrases. This pattern indicates a simple hierarchical rhythmic organisation with a 4:1 ratio between metrical levels. The bird’s behaviour emerged without training, apparent reinforcement, or known exposure to conspecific or human drumming models, suggesting an intrinsic capacity for rhythmic tool use. Although the function of the behaviour remains unclear—play, nutrient extraction, or communicative signalling—these observations extend known rhythmic and tool-using abilities within cockatoos and raise new evolutionary questions. Our findings highlight the potential for rhythmically structured, instrumental behaviour to arise in a broader range of avian taxa than previously recognised, motivating further comparative and experimental work on the cognitive and biomechanical foundations of drumming in parrots.

## Introduction

> *“You’ve got two empty halves of coconut and you’re banging them together!” – Guard (Monty Python and the Holy Grail)*

Music and dance are ubiquitous across human cultures (Mehr et al., 2019), but rare in other species. Although regularly seen as a product of human culture, increasing academic interest has been devoted to understanding musicality—the capacity to create and appreciate music—as an evolved, biological trait (Cross & Morley, 2009; Honing et al., 2015). To understand the evolutionary origins of musicality in humans, both cross-species comparison and archaeological evidence are regularly employed to determine when and why the different component abilities within musicality may have emerged (Fitch, 2015).

The first evidence of tool use for specifically musical purposes comes from bone flutes constructed approximately 40 000 years ago (Montagu, 2017), however, instruments such as bone flutes were almost certainly preceded by rhythmic body-percussion and vocalisations (Morley, 2013). Rhythmic percussion appears to be ancient and widespread across human societies, with certain consistent features such as regular, temporally-organised patterns and isochrony (Fitch, 2012; Ravignani & Madison, 2017). The presence of tool-use for musical purposes, and for drumming in particular, is an important area of investigation to understand the origins of this aspect of musicality.

Drumming is a form of non-vocal communication in which sounds are produced by the impact of either a body part or an object on a substrate (Dunlop et al., 2022; Silberstein et al., 2024). Various forms of drumming exist in non-human animals. Kangaroos (Randall, 2001) and Mongolian Gerbils (*Mongolian gerbil*) will stamp their feet (Silberstein et al., 2024), while Chimpanzees (*Pan troglodytes*) will strike a tree buttress with their hands (Eleuteri et al., 2025). Certain primates, such as Mountain Gorillas (*Gorilla beringei beringei*) will engage with body percussion, seemingly to display size and strength (Wright et al., 2021). Crucially, Chimpanzee and Bonobo (*Pan paniscus*) drumming behaviours exhibit signs of rhythmic structure and non-random timing (Eleuteri et al., 2025).

Tool-assisted drumming is less common than body percussion. The nut-cracking behaviours of Chimpanzees could be considered a form of drumming, although it is usually thought to be primarily for resource extraction purposes (Berdugo et al., 2025). There are also rare instances of Chimpanzees rhythmically throwing stones against tree buttresses, which could be serving a communicative function (van Loon et al., 2025). Aside from mammals, there is also evidence of drumming in birds, which provides valuable insights into the evolution of rhythm processing.

Drumming is uncommon in birds but has been observed in some species. Examples include: the male Ruffed Grouse (*Bonasa umbellus*), which beats its wings to advertise its territory and attract mates (Garcia et al., 2012); manakins (Pipridae), that use multiple mechanisms for producing loud snaps of their wings either against their bodies, or against the air, for courtship purposes (Bostwick & Prum, 2003); and Java Sparrows (*Padda oryzivora*), which use percussive bill-clicks in combination with song (Soma & Mori, 2015). Woodpeckers are likely the most famous avian percussionists, and will rapidly hammer their bills on resonant substrates in territorial and mate-advertising contexts (Brewster, 1876). Although mostly isochronous, their drumming varies between individuals and species in terms of speed, length, and rhythm; these components appear shaped by sexual selection but constrained by biomechanical limits (Schuppe et al., 2021). Brain regions that are associated with drumming in woodpeckers overlap with regions responsible for vocal learning, suggesting convergence in neural circuitry for fine motor communication (Schuppe et al., 2022). That the abilities for fine motor control in both non-vocal and vocal communication may overlap is further supported by the ability of Java Sparrows to coordinate with precise timing across modalities during click and song performance (Soma & Mori, 2015). Although these examples do show evidence of drumming ability across a wide array of birds, they are limited to drumming using the body.

Palm Cockatoos (*Probosciger aterrimus*) present a singular case of instrumental drumming in a bird: in at least two populations, males manufacture and use tools—trimmed sticks or seed pods—to drum on hollow wood during courtship, producing long sequences with isochronous timing and idiosyncratic tempi (Heinsohn et al., 2017). This is the only previously documented case of a non-human animal creating a bespoke tool specifically for sound production. Males show strong, repeatable preferences for tool type and shape, implying aesthetic or functional design choices that may be evaluated by females. These choices may use acoustics to signal information about the size and shape of a potential nesting hollow chosen by the male, during a multimodal courtship performance, which also includes visual displays and vocalisations. Furthermore, there is population-level variation in the prevalence of this behaviour, as only two populations of Palm Cockatoos have been observed drumming, and there appear to be regional dialects between them, hinting at cultural or ecological influences on the emergence and maintenance of drumming traditions in Palm Cockatoos (Heinsohn et al., 2017).

Evidence for rhythmic abilities has been seen in other cockatoo species. The most famous example is Snowball, a dancing Sulphur-crested Cockatoo (*Cacatua galerita eleonora*) that first demonstrated sensorimotor synchronisation in a non-human animal by adapting his movements to the timing of a musical pulse (Patel et al., 2009). Crucially, Snowball also exhibited a wide range of dance movements, demonstrating some creative interpretation (Keehn et al., 2019). Although Snowball has not been observed using tools as part of these rhythmic displays, Sulphur-crested Cockatoos are also adept vocal learners, which has led to speculation that rhythmic coordination is related to vocal learning (Patel et al., 2009), further supported by the aforementioned work on woodpecker neuroanatomy. The musicality and rhythmic ability of Palm Cockatoos, and at least one Sulphur-crested Cockatoo, are noteworthy and raise questions about whether these abilities are widespread amongst cockatoos, or whether these are isolated examples.

The subject of the present study is a Galah (*Eolophus roseicapilla*). Galahs are widespread, Australian cockatoos, and proficient vocal learners. A detailed account of the species is provided in the Handbook of Australian and New Zealand Birds (BirdLife Australia, 2023). In a description of their behavioural ecology, Rowley (1990) identified nine distinct Galah vocalisations. Some Galahs may also dance to human music when prompted (Lubke et al., 2025). No other musical abilities have been previously noted in Galahs. Here we report an instance of a Galah engaging in tool-assisted drumming.

## Method

### Study subject

The subject is a 6.5-year-old, male Galah. The following is a description of him according to STRANGE framework (Webster & Rutz, 2020).

The subject is a captive bird, kept in solitary housing, in a suburban backyard flight aviary, which self-selected to play with items provided for enrichment. It is of captive stock, likely mixed western and eastern sub-species.

The enclosure is adjacent to the rear door of the house so the subject has regular interaction with humans, dogs, wild doves and wild Galahs. Wild Galahs often feed adjacent to the subject, outside the enclosure. The bird communicates by call with its conspecifics using both contact calls and alarm calls to signal the presence of a bird-of-prey. It also mimics some human words; calling the dog, coughing, sneezing, saying ‘hello’ and ‘what are you doing?’.

As a pet bird, the subject interacts with humans, pet dogs as well as visiting doves and wild Galahs which often feed adjacent to the subject, outside the enclosure. It was initially raised by its parents and then moved to its current location from fledging age. It is provided with commercial parrot mix, as well as seeding grass, twigs, nuts, natural branches, perches, logs and commercial parrot toys for enrichment.

The subject began drumming spontaneously at 5.5 years old. As far as the authors are aware, the subject had no prior exposure to drumming as the humans and wild birds with which he has interacted have not demonstrated the behaviour to him. Drumming behaviour occurs at any time during daylight hours, although morning 0830 – 1300 hrs and late afternoon 1630 – 1730hrs seem to be popular times, possibly corresponding to the activity levels of the bird. To date, no seasonality has been observed in the behaviour.

### Field observations and video recordings

Recordings were made using a Sony Handycam HDR-XR520 video camera mounted on a tripod. The camera was set to record and then the operator either came inside so was unable to be seen by the bird or sat reading within the birds’ vision but not looking at or interacting with it. The behaviour was initiated by the bird irrespective of the presence or absence of the operator, provided that the operator did not look at or interact with the bird. The subject was recorded in 14 sessions, totalling approximately 266 minutes, over the course of eight weeks.

### Analysis

Behavioural coding was done using BORIS (Friard & Gamba, 2016). Beat onset—the point at which the coconut impacted the water bowl—was manually coded in one video (Session 8), selected at random. The coder identified a beat on each frame that contained a direct impact between the coconut shell and the metal bowl. If there was any doubt about the precise frame, then the beat would be marked in the frame with that contained the audio peak.

All other sessions were analysed computationally. The audio was separated from the video, then audio peaks were identified using onset detection in Librosa (McFee et al., 2023) and converted to inter-onset intervals (IOIs). Manual observations indicated that an IOI greater than 1.5s indicated a boundary between phrases of drumming, and so a low-pass filter at 1.5s was applied. Kernal density estimation (KDE) and peak estimation were done using SciPy (Virtanen et al., 2020).

## Results

Drumming was first observed in March 2025 (12 months prior, at time of writing), using a half coconut shell. The half shell was part of a commercial parrot toy which was provided to the bird as part of behavioural enrichment. The bird had separated the half coconut shell from the rest of the toy and began manipulating it using a hole which had been drilled in the top. The coconut shell was often carried around or dropped into the bird’s water bowls.

In apparent play behaviour, the bird often deposited toys, leaves, twigs and other items in his water bowls, as well as upending them. One of these water bowls was a metal dog bowl which was no longer used as a water receptacle due to it being regularly upended by the bird. It remained in the enclosure however as the subject seemed to enjoy playing with it. A selection of objects was available for play, however the half coconut and the metal dog bowl were particularly favoured by the bird. They were both turned over and made noise as they knocked into other structures in the enclosure.

It is not known how the subject commenced drumming, however rhythmic behaviour using the coconut shell was first observed approximately 12 months ago.

A rhythmic tapping sound was heard from the enclosure and, after observing the bird covertly, it was determined that the sound was being created by the bird tapping the coconut on the upended metal water bowl. It was difficult to observe the behaviour as the bird would stop if a human observer appeared. Drumming behaviour was never initiated when an observer was present and ceased if the bird perceived that a human was watching it. The bird has been observed to tap with other items; however, the coconut shell and the metal bowl are the only objects which the bird uses for more than a few seconds.

Out of the 14 sessions recorded, Session 8 was randomly chosen for manual analysis. This session contained one main bout of drumming, which lasted 317 seconds and contained 184 beats (Figure 2). Two distinct tapping rates were identified: one with an inter-onset interval of 0.8s and another at 0.2s (see Figure 3, left). These were often arranged into phrases, starting with the longer beats, and then ending with a sequence of short beats, before taking a pause. This indicates a rhythmic structure at two metrical levels, the fast section being 4-times faster than the slow section. Audio analysis of all 14 sessions confirmed the bimodal distribution of inter-onset intervals (Figure 3, right). In total, the subject spent approximately 7.8% of the recorded time actively engaging in drumming behaviour.

**Figure 1.**
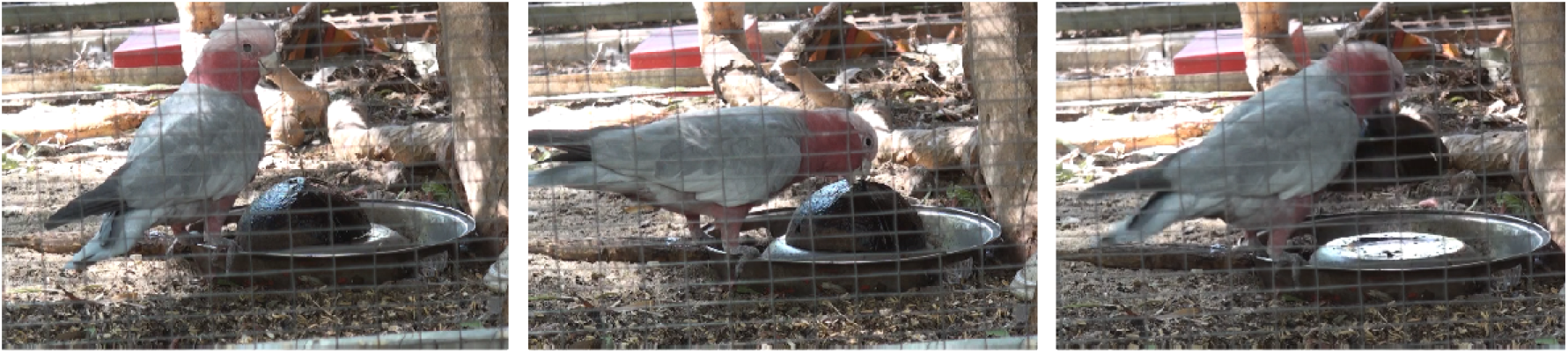
The subject and his half coconut shell and metal dog bowl.

**Figure 2.**
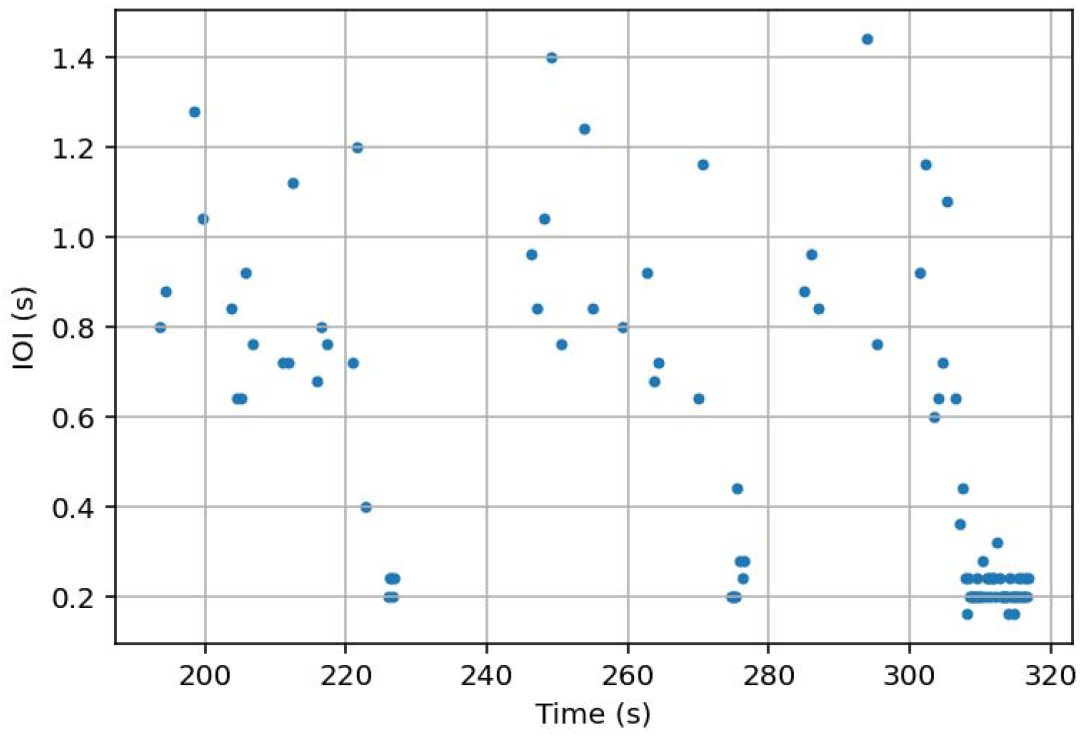
Inter-Onset Intervals over time, identified manually in a segment of Session 8, showing three typical drumming phrases that consist of a set of long beats, followed by a set of short beats in succession, then a pause.

**Figure 3.**
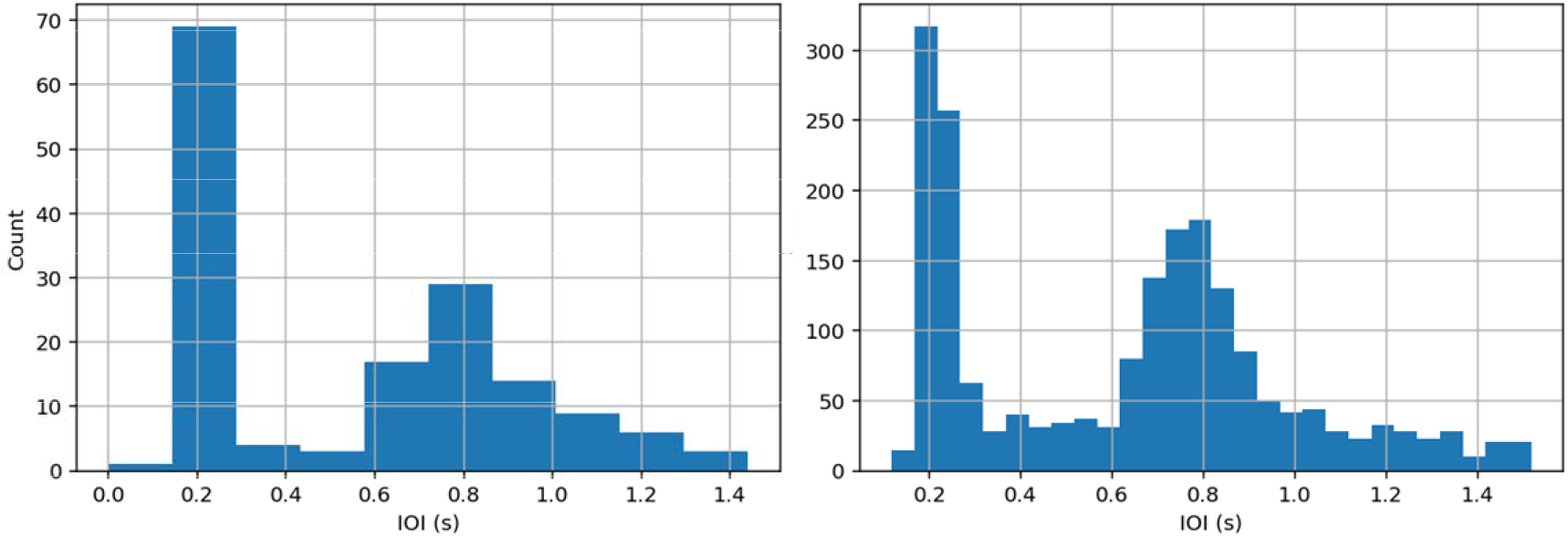
Histograms of Inter-Onset Intervals, identified manually from session 8 (left) and extracted computationally across all recordings (right).

The peaks were confirmed using kernel density estimation (KDE) to obtain a smooth, continuous approximation of the underlying distribution. Local peaks were identified at 0.226 and 0.763, as points where the KDE curve exhibited positive-to-negative derivative changes and exceeded a minimum prominence threshold (0.001). Only direct impacts were included in the manual analysis, while the computational peak extraction includes other audio events. In particular, the subject sometimes would scrape the coconut shell on the water bowl as he picked it up, which made a clear sound preceding the main beat, and this could account for the slightly skewed distribution towards the left of the 0.8 peak in the computational analysis.

## Discussion

Prior to these observations, the literature on instrumental, tool-assisted drumming was limited to certain primates (including Humans), and to the Palm Cockatoo. Here we have demonstrated that at least one other species of cockatoo is capable of drumming using a tool against a substrate.

It must be noted that this behaviour has only been observed in a single individual living in captivity. As such, it represents a capability, rather than proclivity. Drumming has not been observed in wild Galahs, although their abilities to manipulate objects using their bills— rather than their feet, as is common with other parrots—have been previously noted (BirdLife Australia, 2023). While Palm Cockatoos will drum using sticks held in their feet (Heinsohn et al., 2017), the subject here exclusively used his beak, which may be more typical of his species.

Questions remain as to the purpose of drumming in this specific individual. It could have originated as a foraging behaviour, if it were able to extract some nutrients from the coconut shell through repeated impacts. This would make the observed behaviour more analogous to nut-cracking for resource extraction in Chimpanzees (Berdugo et al., 2025), rather than tool-assisted drumming for communicative purposes, as is the case with Palm Cockatoos (Heinsohn et al., 2017). Although the coconut shell was old and dried-up at the time of filming, the subject was observed dipping the shell in water, and licking around the grip-point, so it is possible that he was extracting something of nutritional value.

Another possibility is that the bird is signalling to humans, as it lives in close proximity to humans, and regularly interacts with them. However, the drumming behaviour usually ceases when a human appears or looks at the bird. This could either be because the subject has achieved its aim of attracting a human and therefore no longer needs to drum, or because the drumming is purely for its own purpose and is not something to be shared with humans. Further study may be able to distinguish between these goals. Crucially, the subject does not seem to drum for other Galahs, but does interact with them in other ways, primarily through vocalisations such as standard alarm calls.

Finally, the behaviour could be a form of play. Rowley (1990) observed acrobatics in Galahs which he describes as being like ‘play’ behaviour. They swing from branches and wires, sometimes just by their beaks. He comments that it could appear like ‘boredom’ but seems more like play. The subject of this study spends time manipulating many of the objects available to him however, on a recent occasion, after partially stripping a fresh tree branch provided, preferentially chose drumming as an activity.

Some insights into how the subject learnt to drum may be gained from comparison. Heinsohn and others (Heinsohn et al., 2017) suggest several theories to explain the origin of drumming in Palm cockatoos. Two theories suggest that the birds may have learnt to drum from humans, either by observing and copying drumming by First Peoples of the area, as drumming has been an important ceremonial activity for up to 60,000 years or, more recently, when the birds came into contact with the rock music of humans in the 1960s and 70s. They observe that the Rolling Stones and other music of that era were almost always around a tempo of 120 beats per minute, which is still a highly prevalent tempo in human popular music (Moelants, 2002), and which is similar to the rate that Palm Cockatoos prefer. Therefore, it is possible that the subject of this study could have listened to music played by the humans around him and copied the rhythm from them.

Exactly why and how the subject learnt to drum remains unclear but, nevertheless, the rhythmic structure of the subject’s drumming appears robust. The subject has two clearly defined tapping speeds: a slow tap with a period of approximately 0.8 seconds, and a fast tap with a period of approximately 0.2 seconds. This implies two metrical levels with a simple 4:1 relationship, although it should be made clear that this is not a perfect relationship in our computational analysis. Bowling and Purves (2015) have previously noted the prevalence of simple-integer ratios in animal communication. Such simple ratios are certainly present in human music, although there is usually a preference for 2:1 rhythmic structures across human cultures (Jacoby et al., 2024). Crucially, the rhythmic pattern shown by the subject here could be considered to be isochronous, albeit at two, alternating metrical levels, and isochrony is an important foundation of rhythm (Merker et al., 2009).

The origin of this 4:1 rhythmic structure in the subject remains unclear. On one hand, it could result purely from biomechanical constraints. Even human music is constrained by the range of motion and timing that can be produced by a human body (Godøy, 2018), which may have influenced human preferences for simple ratio rhythmic structures throughout early development (Larsson et al., 2019). In this way, the two speeds seen in this Galah may be similar to the different modes of gait in a horse (Farley & Taylor, 1991). On the other hand, there could be a perceptual preference for this 4:1 ratio, perhaps due to neural architecture.

It was previously suggested that vocal learning is connected with rhythm perception and production. Studies of woodpeckers show that the relevant neural architecture is common to both functions (Schuppe et al., 2022), and Snowball’s dance abilities have been attributed to the vocal learning capabilities of his species (Patel et al., 2009). It is perhaps worth noting that our subject’s fast beat does resemble the very fast and consistent tapping of a woodpecker. This connection raises the possibility that any bird that has both vocal learning, and is adept at object manipulation, could potentially learn to drum. Further comparative studies would be required to confirm if there are other pre-requisite abilities.

The present study is strictly observational, and raises many questions to be investigated experimentally. The subject has only displayed consistent drumming behaviour with one specific coconut shell, so further studies could investigate whether he will drum with any other objects, if provided. The weight of the existing coconut shell could also be manipulated, which may influence the preferred tapping rate if, as suggested earlier, the tapping rate is a function of bio-mechanical constraints. Alternatively, an external pulse could be provided, such as through a musical stimulus, to investigate whether the subjects will beat match. Finally, and although there has, so far, been no indication that the subject is drumming for the wild Galahs that live near him, it could be investigated whether social transmission of the behaviour is possible.

Previous comparative literature into the evolutionary origins of musicality has suggested that human rhythmic abilities are part of a gradual evolution of drumming in primates, while human pitch perception and production represent convergent evolution with birdsong (Honing, 2026). This has perhaps led to a perception that primates do rhythm, while birds do pitch. However, there are now numerous studies of birds—such as Snowball (Patel et al., 2009), the Palm Cockatoos (Heinsohn et al., 2017), and the Galah studied here—that would challenge this view. The precise motivations of the subject remain opaque, however; it is unknown whether this Galah is drumming for food, communication, or play. These motivations may also blur into one another, so that what began as a food source may have become a game. Regardless, these observations suggest that the basic foundations of rhythm exist across a broader range of avian taxa than previously thought. At the very least, many living parrots may be capable of taking a water bowl and half a coconut, and banging them together.

## Ethics

The University of Jyväskylä adheres to the guidelines set out by the Finnish National Board on Research Integrity (TENK) and the Finnish Food Authority as the regulatory body for animal experiments. As this study is strictly observational, it did not require ethics approval, according to the aforementioned guidelines.

## Data accessibility

All data used to generate the results and scripts used in analysis are available from the OSF, along with a video example. https://osf.io/bqx8u/overview?view_only=2ff879ec005149e698f79d6506c57a58

## Declaration of AI use

AI was used to assist with background literature searches and for the generation of code used in data processing. Any code produced in this way was edited by a human author. No AI was used in the writing of this manuscript.

## Authors’ contributions

J.S.B.: conceptualisation, formal analysis, methodology, visualisation, writing—original draft, writing—review and editing; A.R.B.: conceptualisation, investigation, methodology, writing— original draft, writing—review and editing.

## Conflict of interest declaration

We declare that we have no competing interests.

## Funding

This research was funded by the Research Council of Finland (346210) through the Centre of Excellence in Music, Mind, Body and Brain.

## Acknowledgements

The authors would like to acknowledge that this research was conducted on Whadjuk Noongar country, and pay our respects to Elders past and present.

## Notes

https://osf.io/bqx8u/overview?view_only=2ff879ec005149e698f79d6506c57a58

### Competing Interest Statement

The authors have declared no competing interest.

